# Trust, but Verify: Epistemic Uncertainty-Gated Pseudo-Labeling for Safer Semi-Supervised Malaria Screening

**DOI:** 10.64898/2026.10.13.758712

**Authors:** Aman Kumar, Ayush Anand, Latchan Chhetri

**Affiliations:** Halo Mind Research Group, India; Sikkim Manipal Institute of Technology, India

**Keywords:** semi-supervised learning, malaria screening, pseudo-labeling, Monte Carlo dropout, uncertainty quantification, medical image classification, deployment safety, telemedicine, low-resource healthcare

## Abstract

**Background:** Deep learning could make malaria microscopy more accessible in low-resource settings, but training it usually needs thousands of expert-labeled images. Semi-supervised learning (SSL) reduces this need by also learning from unlabeled images. However, methods such as FixMatch accept pseudo-labels using a fixed confidence threshold, and lowering that threshold to use more data can teach the model to make confident mistakes. These silent failures are dangerous because they are never flagged for expert review.

**Methods:** We added an uncertainty check to SSL pseudo-labeling. Each unlabeled image is passed through the model ten times with Monte Carlo dropout active; its pseudo-label is used only if the normalized predictive entropy of the averaged prediction is below 0.2, and it is then weighted by a sigmoid function of model confidence. We trained a ResNet-18 on the public Malaria Cell Images dataset (27,558 cell images; 70:10:20 split) using labels for only 20% of the training images. Over six random seeds, we compared the method with a supervised baseline and with FixMatch at thresholds of 0.95 and 0.80, measuring accuracy, AUC and silent failure rate (SFR), the share of predictions that are wrong despite confidence above 0.95.

**Results:** Compared with FixMatch at a threshold of 0.80, our method lowered the mean SFR from 2.28% to 1.52%, a 33.2% relative reduction (paired t-test, *t* = 4.29, *p* = 0.0078), and the SFR was lower in all six runs. Accuracy (96.6% vs. 96.4%) and AUC (0.991 for both) were unchanged. The mean SFR was also below that of FixMatch at 0.95 (1.97%) and of the supervised baseline (2.01%).

**Conclusions:** Checking epistemic uncertainty before accepting pseudo-labels reduced confident errors in semi-supervised malaria cell classification without lowering accuracy. Patient-level and external validation on whole-slide images are still needed before clinical use.

## 1 Introduction

Malaria is one of the major health issues across the world, with 241 million cases documented in 2020, mostly in resource-limited regions [1]. Microscopic examination of blood smears and detection of parasites are carried out by technicians, which might not always be feasible in resource-limited regions. Deep learning-based approaches have shown promise in automated detection of malaria [2, 3]. However, they are mostly based on thousands of annotated images and samples, which might incur significant costs in such regions.

Another alternative is semi-supervised learning (SSL), which makes use of unlabeled images and samples [4, 5]. However, there are certain risks involved in using SSL-based approaches for deployment in such scenarios. In SSL-based approaches, a certain threshold value is used to choose samples from unlabeled images and samples. Although high threshold values guarantee precision, they are not efficient in utilizing unlabeled images and samples. Lower threshold values are chosen to increase learning efficiency but might incur incorrect and overly confident predictions, which might have severe consequences in such scenarios where human intervention is least involved.

The challenge of threshold selection reveals a fundamental tension in medical SSL deployment. Clinical settings demand both efficiency (to justify automation costs) and safety (to prevent misdiagnosis). Standard confidence-based filtering forces a binary choice between these objectives, lacking mechanisms to distinguish genuinely confident predictions from overconfident errors. This brittleness becomes critical in low-resource contexts where the system may process cases beyond its training distribution without expert verification.

To tackle this problem in the deployment process, we use the method of uncertainty-guided pseudo-label filtering. Instead of relying solely on the model’s confidence level, we use Monte Carlo dropout sampling [6] to estimate the model’s epistemic uncertainty. This way, we can identify the pseudo-labels for which the model is uncertain.

The main contributions of this work are as follows: First, we show the efficacy of integrating uncertainty estimation with SSL by achieving statistically significant improvement in deployment safety for medical imaging (33.2% reduction in silent failure rate, p = 0.0078, N = 6 seeds). Second, we show that the safety improvement is consistent across all six independent training runs. Third, we extend the applicability to malaria screening in low-resource settings and show the efficacy of the method using only 20% of the training labels.

## 2 Related Work

### 2.1 Semi-Supervised Learning

Semi-supervised learning utilizes unlabeled samples for improving the performance of a model when labeled instances are not sufficient. State-of-the-art models utilize consistency regularization [5, 7] and pseudo-labeling [4, 8]. FixMatch [4] is a recent method combining both consistency regularization and pseudo-labeling. The method achieves strong performance across computer vision benchmarks but relies on fixed confidence thresholds that may not transfer to medical domains where class distributions differ and errors carry clinical consequences.

MixMatch [9] and ReMixMatch [10] build upon the distribution alignment methods but share the same limitation due to the use of filtering techniques based on thresholds. These methods assume prediction confidence accurately reflects correctness probability, an assumption that breaks down in medical imaging where models exhibit overconfidence on ambiguous or out-of-distribution samples [11, 12].

### 2.2 Medical Image Classification

Deep learning methods have shown promise in malaria detection [2, 3], tuberculosis screening [13], and other medical diagnosis problems [14, 15], including brain tumor classification and segmentation from MRI [16–18]. However, these successful applications usually require large amounts of labeled data. Semi-supervised deep learning methods have not received much attention in medical imaging problems but can play an important role in such scenarios. Recent work on self-supervised pretraining [19] reduces annotation needs but still requires substantial labeled data for downstream tasks. Evaluation protocols matter as much as model design. Medical image datasets often contain several images per patient, and random image-level splitting can place images of the same patient in both training and test sets, allowing a model to exploit patient-specific features. In our prior work on brain tumor MRI classification, accuracy was 1.23–3.71 percentage points lower under patient-level than under image-level splitting across the evaluated architectures [16], and a controlled ablation under strict patient-level splitting showed that preprocessing and augmentation choices interact and can reduce accuracy [17]. In brain tumor segmentation, we separately isolated the contributions of network architecture and input representation [18]. Like most of the literature, these studies evaluated models with accuracy-, AUC- or overlap-based metrics, which do not distinguish confident errors from uncertain ones; the present work addresses this gap.

There are various challenges in deploying medical AI systems, which are not usually encountered in the performance metrics of the model. Calibration and reliability are important factors in medical AI systems, especially in clinical decision-making problems [11, 20]. Models must not only achieve high accuracy but also provide trustworthy confidence estimates and mechanisms to identify uncertain cases requiring human review. Training-time regularisers such as label smoothing, which we previously applied to discourage overconfident predictions in brain tumor classification [16], can soften output probabilities but do not provide a per-sample estimate of model uncertainty.

### 2.3 Uncertainty Quantification

Epistemic uncertainty estimation allows the model to express its confidence in the acquired knowledge. Bayesian neural networks [21] are used to achieve this in a principled manner but at the expense of increased computation. Monte Carlo dropout [6] is another widely used technique that is more feasible and estimates the model’s uncertainty through the application of dropout layers during the forward pass.

Although the application of uncertainty quantification has been explored in the context of medical image analysis [22, 23], its application in the context of integrating with SSL and its deployment in safe mode is still an unexplored area. Existing work focuses on uncertainty for active learning sample selection [24] rather than pseudo-label filtering for safe training.

## 3 Materials and Methods

### 3.1 Dataset Preparation

We use the publicly available Malaria Cell Images dataset [2], which has 27,558 images of cells, of which 13,779 images are parasitized, and 13,779 images are uninfected. The images are resized to a size of 128 × 128 pixels. We use a 70:10:20 random split of the data to train, validate, and test the model. We simulate low-resource conditions by only exposing 20% of the training labels during learning, with remaining samples treated as unlabeled. The split was performed at the level of individual cell images rather than patients; because multiple cells are cropped from each patient’s blood smear, cells from the same patient may appear in more than one partition (see Section 6).

### 3.2 Problem Formulation

We consider a binary classification task with labeled dataset 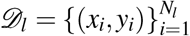 and unlabeled dataset 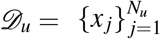 where *N*_*u*_ ≫ *N*_*l*_. Our objective is to train a classifier *f*_*θ*_ : ℝ^*H*×*W* ×3^ → [0, 1]^2^ that minimizes diagnostic errors while maintaining deployment safety.

We define silent failure rate (SFR) as the proportion of samples where the model prediction is both incorrect and highly confident (exceeding threshold *τ*). Formally:

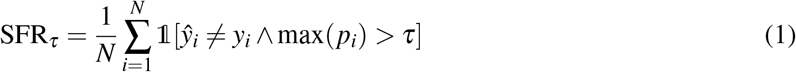

where *p*_*i*_ = *f*_*θ*_ (*x*_*i*_) is the predicted probability distribution. Silent failures represent the most dangerous error mode for deployment, as they bypass uncertainty-based referral mechanisms.

### 3.3 Model Architecture

We employ ResNet-18 [25] pretrained on ImageNet [26] as our backbone network. The final classification layer is replaced with a dropout-enabled head:

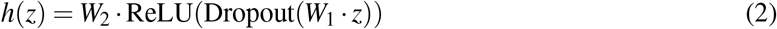

where *z* is the penultimate layer representation and *W*_1_,*W*_2_ are learned weight matrices. Dropout layers (*p* = 0.3) enable uncertainty quantification through stochastic forward passes.

### 3.4 Uncertainty Estimation

Given an input image *x*, we estimate epistemic uncertainty through Monte Carlo sampling. With dropout enabled during inference, we obtain *T* stochastic predictions:

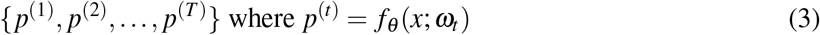

and *ω*_*t*_ represents the *t*-th dropout mask. The mean prediction is:

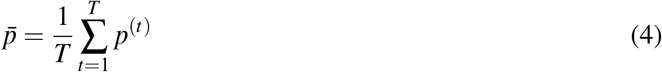

We quantify uncertainty using predictive entropy:

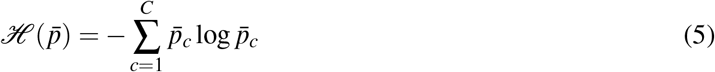

Normalized entropy 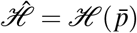 */* log*C* ranges from 0 (certain) to 1 (maximum uncertainty). High entropy indicates model disagreement despite potentially high maximum probability, signaling unreliable predictions.

### 3.5 Adaptive Confidence Weighting

Standard SSL methods apply hard thresholding to select pseudo-labels:

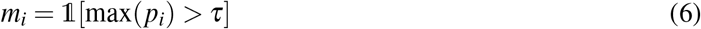

This binary selection discards potentially useful samples with confidence near but below *τ*. We instead employ soft weighting:

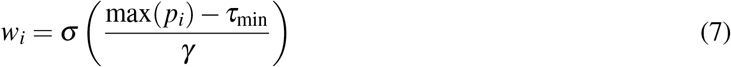

where *σ* is the sigmoid function, *τ*_min_ is minimum acceptable confidence, and *γ* controls transition steepness. This formulation enables gradual incorporation of mid-confidence predictions, improving sample efficiency.

### 3.6 Uncertainty-Guided Training

Figure 1 illustrates our complete training framework. Our training procedure integrates supervised and unsupervised losses. For labeled data, we minimize standard cross-entropy:

**Figure 1.**
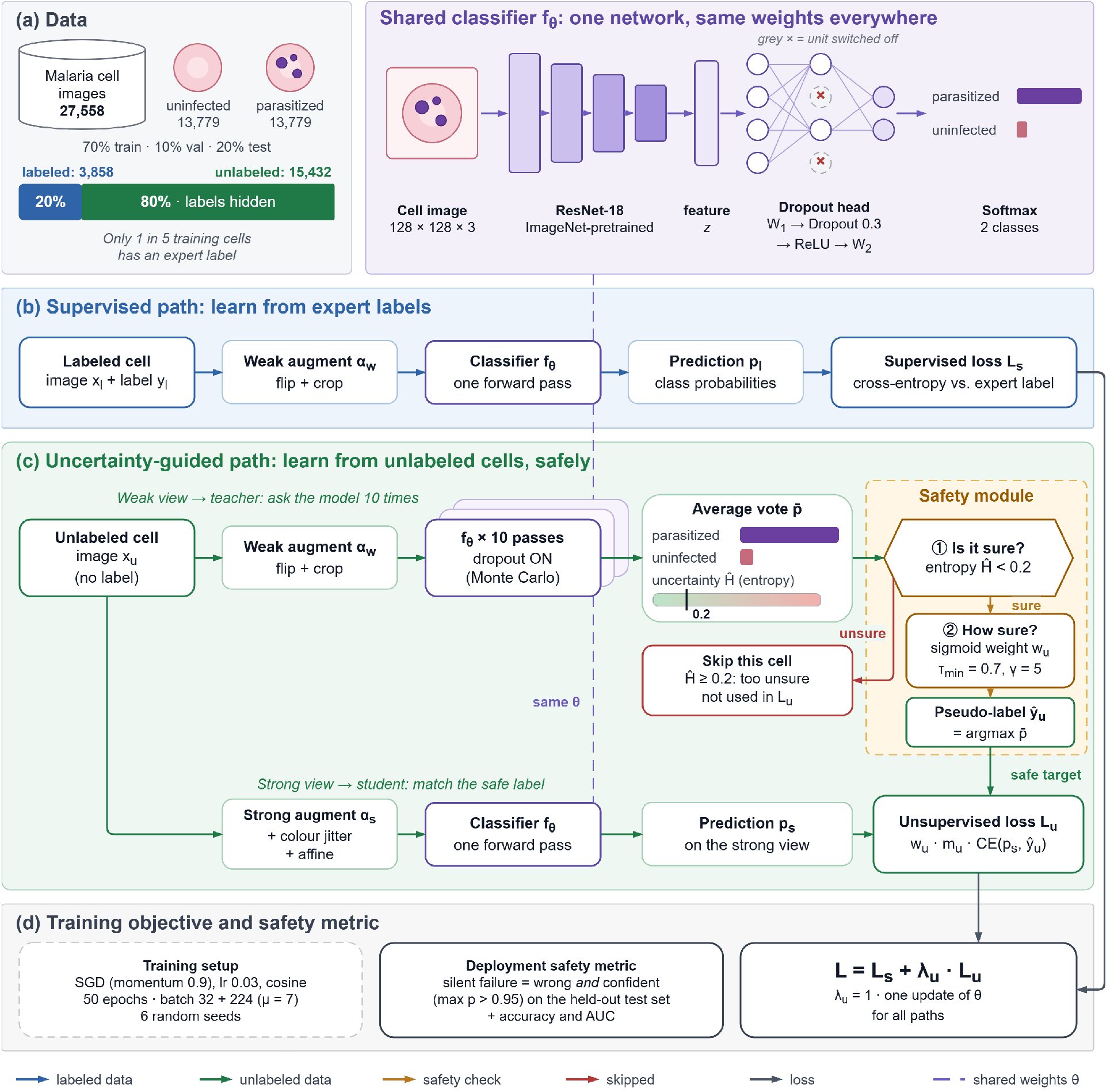
Overview of our uncertainty-guided semi-supervised learning framework. The supervised path (left) processes labeled data through standard cross-entropy loss. The uncertainty-guided path (right) applies Monte Carlo dropout sampling to estimate prediction uncertainty, then uses dual filtering criteria (soft confidence weighting with *τ*_min_ = 0.7, *γ* = 5.0 and normalized entropy *<* 0.2) to ensure safe pseudo-label generation. This two-layer safety mechanism enables efficient learning from unlabeled data while reducing dangerous overconfident errors.

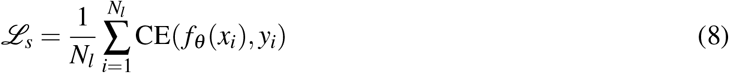

For unlabeled data, we apply consistency regularization with uncertainty-guided filtering. Given unlabeled image *x*_*u*_, we generate weakly augmented version *α*_*w*_(*x*_*u*_) for pseudo-label generation and strongly augmented version *α*_*s*_(*x*_*u*_) for consistency enforcement.

We compute uncertainty-aware weights:

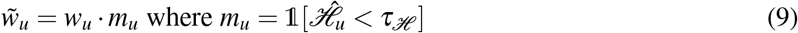

combining confidence weighting *w*_*u*_ with uncertainty filtering *m*_*u*_. The pseudo-label is *ŷ*_*u*_ = argmax 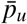 where 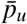 is obtained from Monte Carlo sampling on *α*_*w*_(*x*_*u*_).

The unsupervised loss becomes:

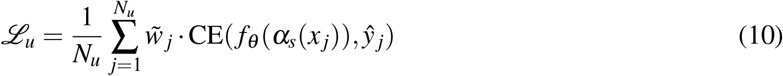

Total training objective:

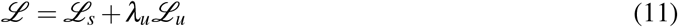

where *λ*_*u*_ balances supervised and unsupervised terms.

### 3.7 Implementation Details

Training is done using the stochastic gradient descent optimizer with momentum of 0.9, weight decay of 5 × 10^−4^, and the initial learning rate of 0.03 with cosine annealing. The batch size is 32 for labeled data and 224 for unlabeled data (*µ* = 7 ratio). The training is done for 50 epochs.

Weak augmentation applies random horizontal flip and crop, while strong augmentation adds color jitter and affine transformations.

For uncertainty estimation, we use *T* = 10 Monte Carlo samples with dropout rate *p* = 0.3. Hyper-parameters are *τ*_min_ = 0.7, *γ* = 5.0, *τ*_*H*_ = 0.2, and *λ*_*u*_ = 1.0. All experiments use PyTorch on NVIDIA GPUs.

### 3.8 Baselines

We compare against:

- **Supervised**: ResNet-18 trained only on 20% labeled data
- **FixMatch (***τ* = 0.95): Standard FixMatch with high confidence threshold
- **FixMatch (***τ* = 0.80): FixMatch with relaxed threshold (tuned baseline)

### 3.9 Evaluation Metrics

Beyond standard accuracy and AUC, we emphasize deployment safety metrics:

- **Silent Failure Rate (SFR)**: Proportion of confident incorrect predictions at threshold *τ* = 0.95
- **Robustness**: Consistency of the safety improvement across independent training runs (random seeds)

All results are averaged over 6 independent training runs with different random seeds. Statistical significance is assessed using paired t-test (*α* = 0.05), comparing our uncertainty-guided method against the tuned baseline (FixMatch *τ* = 0.80) across all seeds. Repeated runs allow the variability of the reported differences to be quantified, which is not possible when each configuration is trained only once [18].

## 4 Results

### 4.1 Main Findings

Table 1 presents our primary experimental results. Our uncertainty-guided framework achieves 1.52% silent failure rate compared to 2.28% for the tuned baseline (FixMatch *τ* = 0.80), representing a 33.2% relative reduction.

**Table 1.** Main Results on Malaria Classification (N = 6 seeds, mean ± std)

| Method | Accuracy | AUC | SFR (%) |
| --- | --- | --- | --- |
| Supervised (20%) | 96.7 ± 0.5 | 0.989 ± 0.004 | 2.01 ± 0.42 |
| FixMatch ($\tau = 0.95$) | 96.8 ± 0.4 | 0.992 ± 0.003 | 1.97 ± 0.38 |
| FixMatch ($\tau = 0.80$) | 96.4 ± 0.5 | 0.991 ± 0.003 | 2.28 ± 0.47 |
| **Ours** | **96.6 ± 0.3** | **0.991 ± 0.002** | **1.52 ± 0.61** |

To check the statistical significance, we ran a paired t-test to compare the silent failure rates across all six independent training runs. The results showed a significant improvement with the paired t-test results showing *t* = 4.29 and *p* = 0.0078 (two-tailed). Therefore, this improvement is significant at the *α* = 0.05 level. The improvement trend is also consistent across all the seeds (Table 2 and Figure 2).

**Table 2.** Silent Failure Rate Per Seed (%)

| Method | Seed 0 | Seed 1 | Seed 2 | Seed 3 | Seed 4 | Seed 5 |
| --- | --- | --- | --- | --- | --- | --- |
| FixMatch | 2.43 | 2.83 | 1.56 | 1.93 | 2.29 | 2.65 |
| Ours | 2.21 | 1.42 | 0.51 | 1.21 | 1.90 | 1.89 |

**Figure 2.**
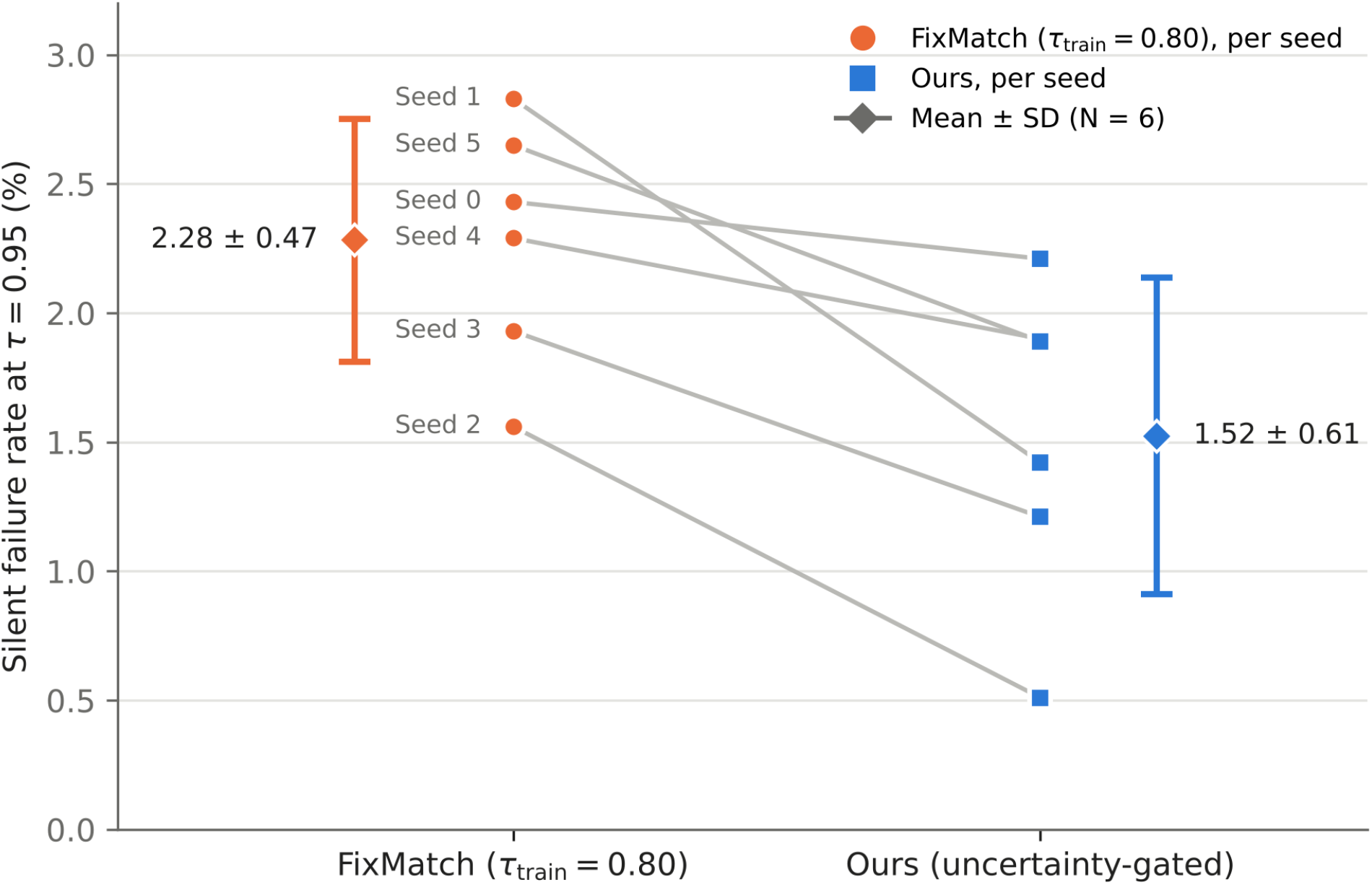
Per-seed silent failure rate at *τ* = 0.95 for the tuned FixMatch baseline (*τ*_train_ = 0.80) and the proposed uncertainty-gated method. Grey lines connect results from the same random seed; diamonds with error bars show the mean ±standard deviation across the six seeds. The markers for seeds 4 and 5 of the proposed method overlap (1.90% and 1.89%). The proposed method has a lower silent failure rate in all six runs (paired t-test, *t* = 4.29, *p* = 0.0078).

Notably, our method also achieves a lower mean silent failure rate than standard FixMatch with the conservative threshold of *τ* = 0.95 (1.52% vs. 1.97%), although this comparison was not tested for statistical significance. The conservative FixMatch configuration limits silent failures by admitting fewer pseudo-labels during training. If the threshold value of the standard FixMatch approach is adjusted to *τ* = 0.80 to use more pseudo-labels, the rate of silent failures rises by approximately 16%.

### 4.2 Per-Seed Analysis

Table 2 and Figure 2 present seed-level results comparing our method to the tuned baseline. Our method has lower silent failure rates than the tuned baseline in all six of the independent trials, with the largest absolute reductions observed on seeds 1 and 2. This consistency across runs suggests that the safety benefit is not an artefact of a particular random initialization.

### 4.3 Clinical Impact

To illustrate the extent to which the proposed improvements can reduce error rates, consider a rural clinic screening 1,000 samples each week. At the observed mean silent failure rates, the baseline algorithm would produce approximately 23 confident misdiagnoses per week, while the proposed algorithm would produce approximately 15, avoiding about 8 silent failures weekly. This corresponds to approximately 394 fewer silent failures annually at a single clinic, assuming a consistent weekly screening volume and error rate. In resource-limited settings where expert review is unavailable, such reduction in silent failures directly impacts patient safety.

## 5 Discussion

### 5.1 Deployment Considerations

Our findings show that the application of uncertainty quantification can lead to the development of safer SSL-based medical imaging systems for use in low-resource settings. The reduction in silent failures by 33.2% addresses the issue of overconfident errors, which is a significant problem in the adoption of automated systems like the proposed solution.

The comparison between the two FixMatch configurations shows the important lesson to be gleaned from the standard SSL methods: the safety level is sensitive to the thresholds used. In the case of deployment scenarios involving varying operating conditions (different imaging hardware, population groups, and disease prevalence), the thresholds might not offer the required safety level. Uncertainty-guided filtering may provide a more robust alternative by explicitly modeling epistemic uncertainty.

### 5.2 Computational Considerations

The computational cost of Monte Carlo dropout sampling is increased by a factor of *T* (which was 10 in our experiments). However, for deployment on resource-constrained devices, the value of *T* could be decreased for screening purposes, while higher values could be reserved for uncertain cases that require review, thus providing a tiered approach for efficiency vs. safety considerations. Alternatively, more recent work on single-pass uncertainty estimation [27] could be leveraged for cost reduction while retaining the filtering benefit.

Another critical aspect for the deployment of the proposed approach for low-resource telemedicine is that, due to the uncertainty-based filtering, not all cases need to be sent for review by experts, thus providing significant savings for the limited-bandwidth connectivity that may exist between remote clinics and the experts for review purposes.

### 5.3 Broader Implications

The problem of deployment safety is not limited to the malaria screening problem but can be generalized to various medical AI applications. As the deployment of automated diagnostic systems increases in environments with limited expert supervision, the need to prevent overconfident errors also arises. Our study shows the potential benefits of integrating uncertainty quantification and semi-supervised learning to solve this problem without losing the benefits of annotation efficiency provided by SSL.

The silent failure rate metric we highlight can be generalized to the broader context of medical AI performance evaluation. In general performance metrics such as accuracy and AUC, the distinction between confident and uncertain errors is not made, and the critical information required to evaluate the deployment safety of AI systems is not provided. As medical AI systems move from the research to the application domain, the need to include deployment safety metrics arises.

## 6 Limitations

Our evaluation is limited to a specific medical imaging task (malaria screening), as well as binary classification. Further validation is necessary across other diagnostic imaging tasks, including multi-class classification, for broader applicability. Although the Malaria Cell Images dataset is suitable for developing the proposed approach, the images provided are crops of cell images, not the entire slide images that might be seen during clinical practice. In addition, the data were partitioned at the cell level rather than the patient level. Because each patient contributes many cells, cells from the same patient may appear in both the training and test sets, which can allow a model to exploit patient- or slide-specific appearance and may inflate absolute performance estimates. In our prior brain tumor MRI study, accuracy was up to 3.71 percentage points lower under patient-level than under image-level splitting [16]. All methods compared here were evaluated under the same protocol, so the comparison between them is controlled, but the absolute accuracy and silent failure rates should be confirmed with a patient-level split.

The proposed uncertainty estimation technique is based on the dropout approximation of Bayesian inference. Although this is efficient, it may not provide the most accurate results, as discussed by Wilson et al. [28]. More advanced uncertainty estimation may also be beneficial for the reliability of the filtering results.

We also did not address the problem of covariate shift between the training and deployment environments, as is common in medical imaging due to the variety of imaging protocols, staining methods, and populations across different sites. Although uncertainty estimation is useful for out-of-distribution detection, more comprehensive solutions for distribution monitoring and adaptation would be necessary for deployment. Finally, silent failure rates were evaluated at a single confidence threshold (*τ* = 0.95). How the advantage of uncertainty gating changes across evaluation thresholds, and across the training thresholds of the baseline, remains to be characterised in future work.

## 7 Conclusion

We proposed an uncertainty-guided semi-supervised learning scheme for medical image classification in resource-constrained environments. By leveraging the effectiveness of the Monte Carlo dropout method to estimate model uncertainty, combined with adaptive confidence weighting, our proposed scheme was found to provide a statistically significant improvement in the safety of model deployments (33.2% reduction in silent failures, p = 0.0078) while retaining the model’s diagnostic performance using only 20% of the training labels. The improvement was also consistent across all six independent training runs, with a lower silent failure rate than the tuned baseline in every run.

In the context of the proposed scheme’s evaluation on the task of malaria cell classification, the work contributes to the development of reliable automated diagnostic tools that may be used to address the problem of underserved regions where expert supervision is scarce. Future work will evaluate the approach under strict patient-level data splitting, as adopted in our prior medical imaging studies [16, 17], and on whole-slide images from external sites.

The road to equitable global health outcomes calls for AI solutions that balance efficiency and safety. Our proposed work shows that uncertainty-guided semi-supervised learning offers promise to bridge this divide, enabling the practical deployment of AI-based diagnostic tools where they may be used to the best effect.

## Declarations

### Ethics statement

This work used only the publicly available, de-identified Malaria Cell Images dataset, released for research use by the U.S. National Library of Medicine [2]. The dataset consists of segmented red blood cell images from Giemsa-stained thin blood smear slides and contains no identifiable patient information. No new data were collected from human participants, and the authors had no access to identifiable information. As the analyses were performed using openly available, de-identified secondary data, no additional participant recruitment or prospective human-subject procedures were undertaken.

### Data availability

The Malaria Cell Images dataset analysed in this study is publicly available, as described in its reference publication [2], and can be obtained from Kaggle at https://www.kaggle.com/datasets/iarunava/cell-images-for-detecting-malaria. No new data were generated. The partitioning procedure (a 70:10:20 random split into training, validation and test sets, with 20% of the training labels exposed during learning) is described in Section 3.1 to facilitate replication. All numerical results reported in this manuscript are contained in Tables 1 and 2 and Figure 2.

### Code availability

To facilitate reproducibility and further research, the code and experimental configurations used in this study are publicly available on GitHub at https://github.com/HaloMind-Research/SafeMed-SSL under the MIT license.

### Author contributions

Aman Kumar, Ayush Anand and Latchan Chhetri contributed equally to this work. All authors contributed to the conceptualisation, methodology, software, formal analysis, investigation, visualisation, and the writing, review and editing of the manuscript. All authors reviewed and approved the final manuscript. Latchan Chhetri is the corresponding author.

## Acknowledgements

The authors sincerely thank Dr. Palash Ghosal for his teaching in biomedical image analysis, his valuable advice, and the insightful discussions that helped shape our understanding and interest in medical imaging.

## Use of generative AI

Generative AI was used to assist with the formatting of this manuscript.

## Reporting guideline

This study was reported with reference to the Checklist for Artificial Intelligence in Medical Imaging (CLAIM 2024) [29].

